# *In silico* investigation of coding variants potentially affecting the functioning of the glutamatergic N-methyl-D-aspartate receptor in schizophrenia

**DOI:** 10.1101/429258

**Authors:** Antonia Tsavou, David Curtis

## Abstract

**Background:** Several lines of evidence support the hypothesis that impaired function of the glutamatergic N-methyl-D-aspartate receptor (NMDAR) might be involved in the aetiology of schizophrenia. NMDAR is activated by phosphorylation by Fyn and there is also some evidence to suggest that abnormalities in Fyn functionality could also be involved in susceptibility to schizophrenia. In a recent weighted burden analysis of exome sequenced schizophrenia cases and controls we noted modest statistical evidence for an enrichment of rare, functional variants in *FYN, GRIN1* and *GRIN2B* in schizophrenia cases.

**Aims:** To test the plausibility of the hypothesis that schizophrenia susceptibility might be associated with genetic variants predicted to cause impaired functioning of NMDAR, either directly or indirectly through impairment of the kinases which phosphorylate it.

**Method:** In an exome sequenced sample of 4225 schizophrenia cases and 5834 controls rare variants occurring in genes for the NMDAR subunits and for the kinases acting on it were annotated. The counts of disruptive and damaging variants were compared between cases and controls and the distribution of amino acids affected by damaging variants were visualised in ProteinPaint and the RCSB Protein Data Bank. Special attention was paid to tyrosine residues subject to phosphorylation.

**Results:** There was no suggestion that abnormalities of the serine-threonine kinases or of Src were associated with schizophrenia. 3 cases and no controls had a disruptive variant in *GRIN2A* and 2 cases and no controls had a disruptive variant in *FYN*. 14 cases and 3 controls had damaging variants in *FYN* and all the variants in controls affected amino acid residues in the N-terminal region outside of any known functional domains. By contrast, 10 variants in cases affected amino acids in functional domains and in the 3D structure of Fyn two of the amino acid substitions, A376T and Q517E, were adjacent to each other. 8 cases and 1 control had damaging variants in *GRIN1* but there was no obvious pattern with respect to particular functional domains being affected in this or other genes. A single case had a variant in *GRIN2A* affecting a well-supported phosphorylation site, Y943C, and three cases had a variant which produce an amino acid change, T216S, which lies two residues away from two adjacent well-supported phosphorylation sites in *FYN*. Aside from this, there was no suggestion that tyrosine phosphorylation sites in Fyn or NMDAR were affected.

**Conclusions:** The numbers of variants involved are too small for firm conclusions to be drawn. The results are consistent with the hypothesis that about 0.5% of subjects with schizophrenia have disruptive or damaging genetic variants which could plausibly impair functioning of NMDAR directly or indirectly through impairing Fyn function.

## Introduction

Outside of the cerebellum, the glutamatergic N-methyl-D-aspartate receptor (NMDAR) consists of a tetramer of two NR1 subunits, coded for by the gene *GRIN1*, and two NR2 units, either NR2A or NR2B, coded for by *GRIN2A* and *GRIN2B* (Chen and Roche, 2007). Variants in these genes can lead to with epilepsy, intellectual disability and other neuropsychiatric disorders (XiangWei, Jiang and Yuan, 2018). As recently reviewed, there is evidence from a variety of sources to support the hypothesis that hypofunction of NMDAR may contribute to the pathophysiology of schizophrenia (Balu, 2016). Phencyclidine, which is an NMDAR channel blocker, induces a psychotomimetic state that closely resembles schizophrenia and can produce not only positive symptoms of hallucinations and delusions but also formal thought disorder and negative symptoms (Javitt and Zukin, 1991). Some studies have found reduced levels of *GRIN1* mRNA and/or NRl protein in different brain regions of subjects with schizophrenia (Gao *et al*., 2000; Law and Deakin, 2001; Stan *et al*., 2015; Catts *et al*., 2016). Patients with autoimmune NMDAR encephalitis can present with psychotic symptoms, memory deficits and catatonic symptoms (Dalmau *et al*., 2011) and patients with an initial diagnosis of schizophrenia are reported to have increased rates of NMDAR antibodies (Steiner *et al*., 2013). *GRIN2A* was included in one of the 108 implicated regions in a large genome-wide association study of schizophrenia (Schizophrenia Working Group of the Psychiatric Genomics Consortium, 2014) and the set of genes coding for proteins associated with NMDAR was found to be enriched for both *de novo* copy number variants (CNVs) and small *de novo* mutations (Kirov *et al*., 2012; Fromer *et al*., 2014). In the Swedish schizophrenia study including nearly 5000 exome-sequenced schizophrenia cases there was an enrichment of ultra-rare, singleton variants predicted to be gene-disruptive among the set of genes encoding interactors with NMDAR (Genovese *et al*., 2016).

A fundamental mechanism for regulating NMDAR function is by phosphorylation, which affects both the trafficking of the receptor and its channel properties, as has been reviewed in detail elsewhere (Chen and Roche, 2007). The subunits contain many serine/threonine phosphorylation sites which are substrates for protein kinases A, B and C (PKA, PKB and PKC), calcium/calmodulin-dependent protein kinase II (CaMKII) and casein kinase II (CKII). Phosphorylation of these sites can have a variety of effects on the pharmacological functioning of NMDAR. Separately, the subunits also contain tyrosine residues which are substrates for phosphorylation by protein tyrosine kinases, in particular Fyn and Src, and this can affect NMDAR function as well as trafficking and endocytosis so that tyrosine phosphorylation by these kinases serves to upregulate NMDAR (Ali and Salter, 2001). The Tyr 1325 residue of NR2A is most prominently phosphorylated *in vitro* by Fyn and mice engineered to have a homozygous Tyr-1325-Phe mutation to prevent the phosphorylation of this site are reported to show some antidepressant-like behaviour (Taniguchi *et al*., 2009). Fyn deficient mice show an increased acoustic startle response under bright light and this is accompanied by increased serotonergic and dopaminergic activity in the prefrontal cortex and limbic system (Hironaka, Yagi and Niki, 2002). Haloperidol injection activates Fyn in mice and increases NR2B phosphorylation while Fyn deficient mice are resistant to haloperidol-induced catalepsy (Hattori *et al*., 2006). Alternative mRNA splicing with lower levels of Fyn protein has been reported to occur in schizophrenia cases and their relatives (Hattori *et al*., 2009) and in a recent GWAS of schizophrenia rs7757969, which lies in an intron of *FYN*, the gene which codes for Fyn, was significant at p = 4.8 × 10^−8^ (Li *et al*., 2017). Of note, Fyn itself is subject to phosphorylation and is activated by autophosphorylation of a conserved residue, tyrosine 416 (Y416), a process which is subject to dopaminergic and cholinergic regulation (Mao and Wang, 2015).

We carried out an analysis of the exome-sequenced Swedish schizophrenia cases and controls but in contrast to the previously reported study (Genovese *et al*., 2016) we did not restrict attention to only singleton variants predicted to completely disrupt gene-functioning but we included all rare variants, though with a weighting scheme to give higher weight to variants predicted to have a more deleterious effect (Curtis *et al*., 2018). For this weighted burden analysis the strength of evidence implicating each gene is summarised as the signed log p value (SLP) and we noted that the set of genes with most evidence for an excess of rare, damaging variants in schizophrenia cases was the set denoted “FMRP targets” (gene set SLP = 7.1) and that within this set the most strongly implicated gene was *FYN*, with SLP = 3.4. Other genes within this set included *GRIN1* (SLP = 1.7) and *GRIN2B* (SLP = 2.1).

These results suggested to us the possibility that genetic variants might increase susceptibility to schizophrenia by impairing the normal functioning of NMDAR through different mechanisms. This might be through variants within the genes coding for the NMDAR subunits or through variants in genes such as *FYN* whose products would normally upregulate NMDAR. Such variants might completely disrupt the functioning of the gene, by producing stop codons, frameshifts or splice site changes. Alternatively they might introduce amino acid changes which critically affected the functioning of the protein. In particular, a coding variant changing a tyrosine residue which was a target of Fyn might have a similar effect to a deleterious variant in *FYN* itself. Accordingly we decided to investigate more closely the coding variants occurring in genes for the NMDAR subunits and the kinases acting on them.

## Methods

We used the results from the previously reported weighted burden analysis of exome-sequenced Swedish subjects consisting of 4225 schizophrenia cases and 5834 controls (Curtis *et al*., 2018). We restricted attention to the genes which code for the subunits of the NMDAR protein and the kinases which are involved in its phosphorylation, as listed in Table 1. Variants were annotated with using VEP, PolyPhen and SIFT (Kumar, Henikoff and Ng, 2009; Adzhubei, Jordan and Sunyaev, 2013; McLaren *et al*., 2016). We restricted attention to variants which were both rare, occurring in a total of no more than 10 subjects, and predicted either to completely disrupt gene functioning (frameshift, stop-gained and essential splice site variants) or to be potentially damaging, defined as being non-synonymous variants predicted as deleterious by SIFT and/or as possibly or probably damaging by PolyPhen. Population frequencies for all variants studied were obtained from gnomAD (Lek *et al*., 2016). For the damaging variants, we visualised the distribution of the altered amino acid residues across functional domains of the protein using the ProteinPaint web application provided at https://pecan.stjude.cloud/home (Zhou *et al*., 2016). In order to assess whether damaging variants might disrupt phosphorylation sites we used the information provided by PhosphoSitePlus (Hornbeck *et al*., 2015). For each protein, this provides information regarding which residues are subject to phosphorylation and other modifications along with the number of references which support this modification from high-throughput and low-throughput experiments, the latter being more reliable. For the assessment of the possible impact of variants on phosphorylation sites we restricted attention to tyrosine residues in the NMDAR subunits and Fyn. In order to visualise the distribution of altered amino acid residues in the 3D structure of the protein we obtained structures from the RCSB Protein Data Bank at www.rcsb.org (Berman *et al*., 2000) and colour-coded altered residues using the JSmol viewer provided there. Since the preliminary results did not suggest that disturbance of serine-threonine kinase functioning was associated with schizophrenia, for the 3D studies we restricted investigation to NMDAR, Fyn and Src.

**Table 1.** Genes for the NMDAR protein and the kinases involved in its phosphorylation.

| **Protein** | **Gene(s)** |
| --- | --- |
| Glutamatergic NMDA receptor (NMDAR) | *GRIN1, GRIN2A, GRIN2B* |
| Protein kinase A (PKA) | *PRKACA, PRKACB, PRKACG, PRKAR1A, PRKAR1B, PRKAR2A, PRKAR2B* |
| Protein kinase C (PKC) | *PRKCA, PRKCB, PRKCG* |
| Protein kinase B (PKB) | *PTK2B* |
| Calcium/calmodulin-dependent protein kinase II (CaMKII) | *CAMK2A, CAMK2B, CAMK2G, CAMK2D* |
| Casein kinase II (CKII) | *CSNK2A1, CSNK2A2, CSNK2B* |
| Proto-oncogene tyrosine-protein kinase Fyn | *FYN* |
| Proto-oncogene tyrosine-protein kinase Src | *SRC* |

## Results

Table 2 shows the summary results for the genes involved as obtained from the original weighted burden analysis. This shows that *GRIN1, GRIN2B* and *FYN* have modestly positive SLP values. In this analysis, three disruptive mutations had been assigned to *GRIN2B*. However on further examination it transpired that there was a single frameshift mutation in one subject but that this had been annotated as three separate frameshift mutations close to each other. Once this was corrected there was one case and one control with a disruptive variant in *GRIN2B* and no excess of damaging variants in cases. 2 cases and no controls had a disruptive variant in *FYN* and both *GRIN1* and *FYN* had an excess of damaging variants in cases. Of note, although *GRIN2A* produced only a very small SLP in the weighted burden analysis (0.14), three cases and no controls had a disruptive variant in this gene. By contrast, *SRC* had an excess of damaging variants in controls and overall there was no suggestion that there was an excess of rare disruptive and damaging variants in cases in the genes coding for the serine-threonine kinases. Table 3 shows a detailed breakdown of the amino acid changes in *GRIN1, GRIN2A, GRIN2B* and *FYN*.

**Table 2.**
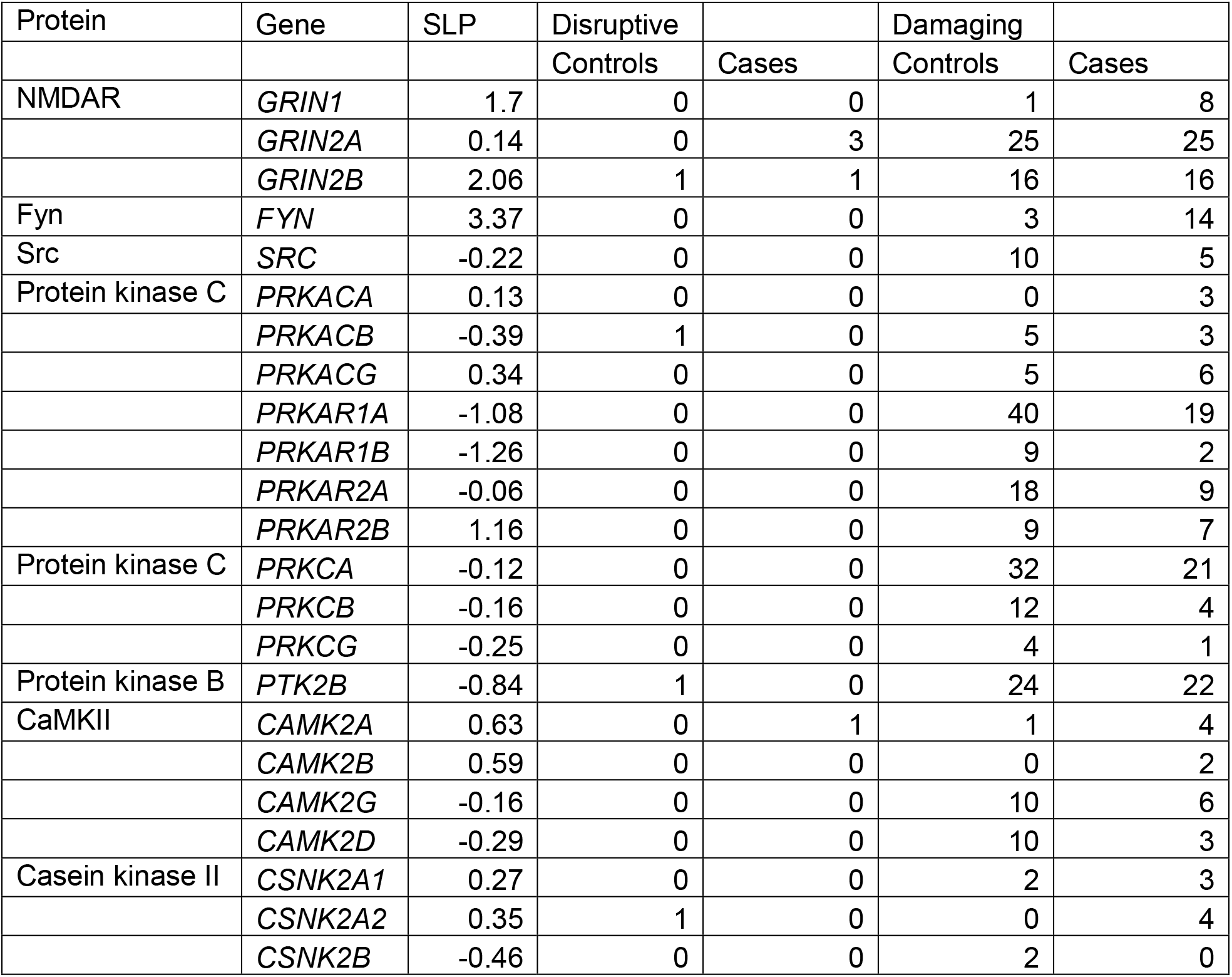
Results obtained from original weighted burden analysis for genes coding for NMDAR and the kinases acting on it. The SLP is the signed log p value, indicating the overall level of support for association with schizophrenia obtained from the weighted burden analysis. For rare variants, occurring in no more than ten subjects, the total counts for gene-disruptive and gene-damaging variants in controls and cases is shown.

**Table 3.** Table showing effects of rare disruptive and damaging variants in *GRIN1, GRIN2A, GRIN2B* and *FYN* along with population allele frequencies obtained from gnomAD.

| Gene | Position | DNA change | Amino acid change as called by VEP | Count in controls | Count in cases | Non-Finnish European allele frequency | Finnish allele frequency |
| --- | --- | --- | --- | --- | --- | --- | --- |
| GRIN1 | 9:140034149 | Gcc/Acc | A/T | 0 | 1 | 9.13E-06 | 4.49E-05 |
|  | 9:140052907 | Gcc/Acc | A/T | 1 | 2 | 0.000179 | 0 |
|  | 9:140052955 | Ggc/Cgc | G/R | 0 | 1 | 8.99E-06 | 0 |
|  | 9:140055795 | aAg/aTg | K/M | 0 | 1 | 9.04E-06 | 0 |
|  | 9:140057671 | tCg/tGg | S/W | 0 | 1 | 8.99E-06 | 0 |
|  | 9:140057749 | cCc/cTc | P/L | 0 | 1 | 8.98E-06 | 0 |
|  | 9:140059705 | aCc/aTc | T/I | 0 | 1 | 8.96E-06 | 0 |
| GRIN2A | 16:9857046 | cGc/cAc | R/H | 0 | 1 | 2.69E-05 | 0 |
|  | 16:9857305 | Cct/Gct | P/A | 1 | 0 | 8.97E-06 | 0 |
|  | 16:9857386 | ggGAaa/ggAaa | GK/GX | 0 | 2 | 2.69E-05 | 0 |
|  | 16:9857503 | Agg/Ggg | R/G | 3 | 3 | 0.000188 | 0 |
|  | 16:9857517 | aTt/aCt | I/T | 1 | 2 | 7.17E-05 | 0 |
|  | 16:9857532 | cAt/cCt | H/P | 0 | 1 | 8.96E-06 | 0 |
|  | 16:9857538 | cGt/cAt | R/H | 1 | 0 | 8.96E-06 | 0 |
|  | 16:9857823 | tTg/tGg | L/W | 3 | 0 | 0.000206 | 0 |
|  | 16:9857833 | Cac/Tac | H/Y | 1 | 1 | 1.79E-05 | 0 |
|  | 16:9857974 | Gag/Aag | E/K | 2 | 0 | 1.79E-05 | 0 |
|  | 16:9858040 | Gat/Tat | D/Y | 0 | 1 | 8.98E-06 | 0 |
|  | 16:9858493 | Cgg/Tgg | R/W | 1 | 0 | 2.69E-05 | 0 |
|  | 16:9858511 | Caa/Aaa | Q/K | 1 | 0 | 8.96E-06 | 0 |
|  | 16:9858573 | tAc/tGc | Y/C | 0 | 1 | 8.96E-06 | 0 |
|  | 16:9858666 | tCc/tGc | S/C | 0 | 2 | 3.58E-05 | 0 |
|  | 16:9858690 | aTt/aCt | I/T | 0 | 1 | 8.96E-06 | 0 |
|  | 16:9858774 | aTt/aCt | I/T | 3 | 1 | 7.19E-05 | 4.48E-05 |
|  | 16:9862746 | Tgc/Cgc | C/R | 0 | 1 | 8.98E-06 | 0 |
|  | 16:9862766 | cGc/cAc | R/H | 1 | 0 | 1.80E-05 | 0 |
|  | 16:9892242 | Atc/Gtc | I/V | 0 | 3 | 2.69E-05 | 0 |
|  | 16:9916247 | cGa/cAa | R/Q | 1 | 0 | 8.96E-06 | 0 |
|  | 16:9928037 | Gtt/Ctt | V/L | 1 | 0 | 8.98E-06 | 0 |
|  | 16:9934519 | Cct/Gct | P/A | 0 | 1 | 0 | 0 |
|  | 16:9934902 | aTt/aGt | I/S | 0 | 1 | 8.97E-06 | 0 |
|  | 16:9943670 | cCc/cAc | P/H | 0 | 1 | 4.48E-05 | 0 |
|  | 16:9984857 | Cgg/Tgg | R/W | 0 | 1 | 8.96E-06 | 0 |
|  | 16:10031933 | gGc/gCc | G/A | 1 | 0 | 2.70E-05 | 0 |
|  | 16:10032093 | Cgc/Tgc | R/C | 0 | 1 | 8.99E-06 | 0 |
|  | 16:10032122 | gAc/gCc | D/A | 1 | 0 | 8.99E-06 | 0 |
|  | 16:10032128 | tCc/tTc | S/F | 1 | 0 | 8.99E-06 | 0 |
|  | 16:10032197 | tCc/tTc | S/F | 2 | 0 | 1.80E-05 | 0 |
|  | 16:10032276 | Ttc/Atc | F/I | 0 | 1 | 9.00E-05 | 0 |
|  | 16:10273938 | Cag/Gag | Q/E | 0 | 1 | 0 | 0 |
|  | 16:10274097 | Gag/Tag | E/* | 0 | 1 | 1.81E-05 | 0 |
| GRIN2B | 12:13715832 | tGt/tAt | C/Y | 0 | 1 | 8.95E-06 | 0 |
|  | 12:13715850 | cGt/cCt | R/P | 1 | 0 | 8.95E-06 | 0 |
|  | 12:13716060 | tAc/tGc | Y/C | 2 | 3 | 7.16E-05 | 4.48E-05 |
|  | 12:13716273 | cGg/cAg | R/Q | 1 | 0 | 8.95E-06 | 0 |
|  | 12:13716318 | cCg/cGg | P/R | 0 | 1 | 8.95E-06 | 0 |
|  | 12:13716424 | Aac/Gac | N/D | 1 | 0 | 8.96E-06 | 0 |
|  | 12:13716667 | Gga/Aga | G/R | 1 | 0 | 1.80E-05 | 0 |
|  | 12:13716940 | Gcc/Tcc | A/S | 1 | 1 | 1.79E-05 | 0 |
|  | 12:13716970 | Gtc/Atc | V/I | 1 | 0 | 1.79E-05 | 0 |
|  | 12:13716998 | tcc/tcAACCc | S/STX | 0 | 1 | 8.95E-06 | 0 |
|  | 12:13717001 | aGc/aTc | S/I | 1 | 0 | 8.95E-06 | 0 |
|  | 12:13717004 | gaC/gaA | D/E | 0 | 1 | 8.95E-06 | 0 |
|  | 12:13717035 | Ggc/Agc | G/S | 0 | 1 | 0.000269 | 0 |
|  | 12:13717061 | cTg/cCg | L/P | 1 | 0 | 8.95E-06 | 0 |
|  | 12:13717096 | aGg/aAg | R/K | 2 | 6 | 0.000134 | 0 |
|  | 12:13717104 | Agc/Ggc | S/G | 1 | 0 | 9.14E-06 | 0 |
|  | 12:13717125 | Gaa/Aaa | E/K | 0 | 1 | 8.97E-06 | 0 |
|  | 12:13717564 | Gtc/Ttc | V/F | 1 | 0 | 8.96E-06 | 0 |
|  | 12:13720189 | tgT/tgA | C/* | 1 | 0 | 0 | 0 |
|  | 12:13764713 | Ctt/Att | L/I | 0 | 1 | 1.81E-05 | 0 |
|  | 12:13768544 | gAt/gTt | D/V | 1 | 0 | 9.03E-06 | 0 |
|  | 12:13828708 | aTc/aCc | I/T | 1 | 0 | 0 | 0 |
|  | 12:13828743 | Cag/Gag | Q/E | 0 | 1 | 9.85E-05 | 0 |
|  | 12:13906758 | Gca/Aca | A/T | 0 | 2 | 6.27E-05 | 0 |
| FYN | 6:111983007 | Tta/Ata | L/I | 0 | 2 | 1.79E-05 | 0 |
|  | 6:112015725 | caG/ca | Q/X | 0 | 1 | 8.95E-06 | 0 |
|  | 6:112015890 | aCc/aGc | T/S | 0 | 3 | 2.69E-05 | 0 |
|  | 6:112021352 | caG/ca | Q/X | 0 | 1 | 8.98E-06 | 0 |
|  | 6:112024138 | cGc/cAc | R/H | 0 | 1 | 2.69E-05 | 0 |
|  | 6:112025201 | T/C | Splice donor variant | 0 | 1 | 8.97E-06 | 0 |
|  | 6:112029200 | Cgt/Tgt | R/C | 1 | 0 | 1.80E-05 | 4.49E-05 |
|  | 6:112029218 | Gtc/Atc | V/I | 1 | 2 | 0.00034 | 0 |
|  | 6:112041026 | tAt/tGt | Y/C | 1 | 0 | 8.96E-06 | 0 |

As shown in Figure 1 and Supplementary Figure 1, for almost all genes the distribution of damaging variants across protein domains did not reveal any particular patterns and there was no marked tendency for a specific functional domain to be enriched for damaging variants in cases. NR1 has an excess of damaging variants in cases and these are widely distributed The only gene to show a pattern in the distribution of damaging variants was Fyn. As shown in Figure 1d and Supplementary Figure 1d, the three amino acid residues altered in controls all occurred in the N-terminal region of the protein to which no functional domain is assigned. By contrast, ten cases had a genetic variant which would alter an amino acid residue in one of the functional domains.

**Figure 1.**
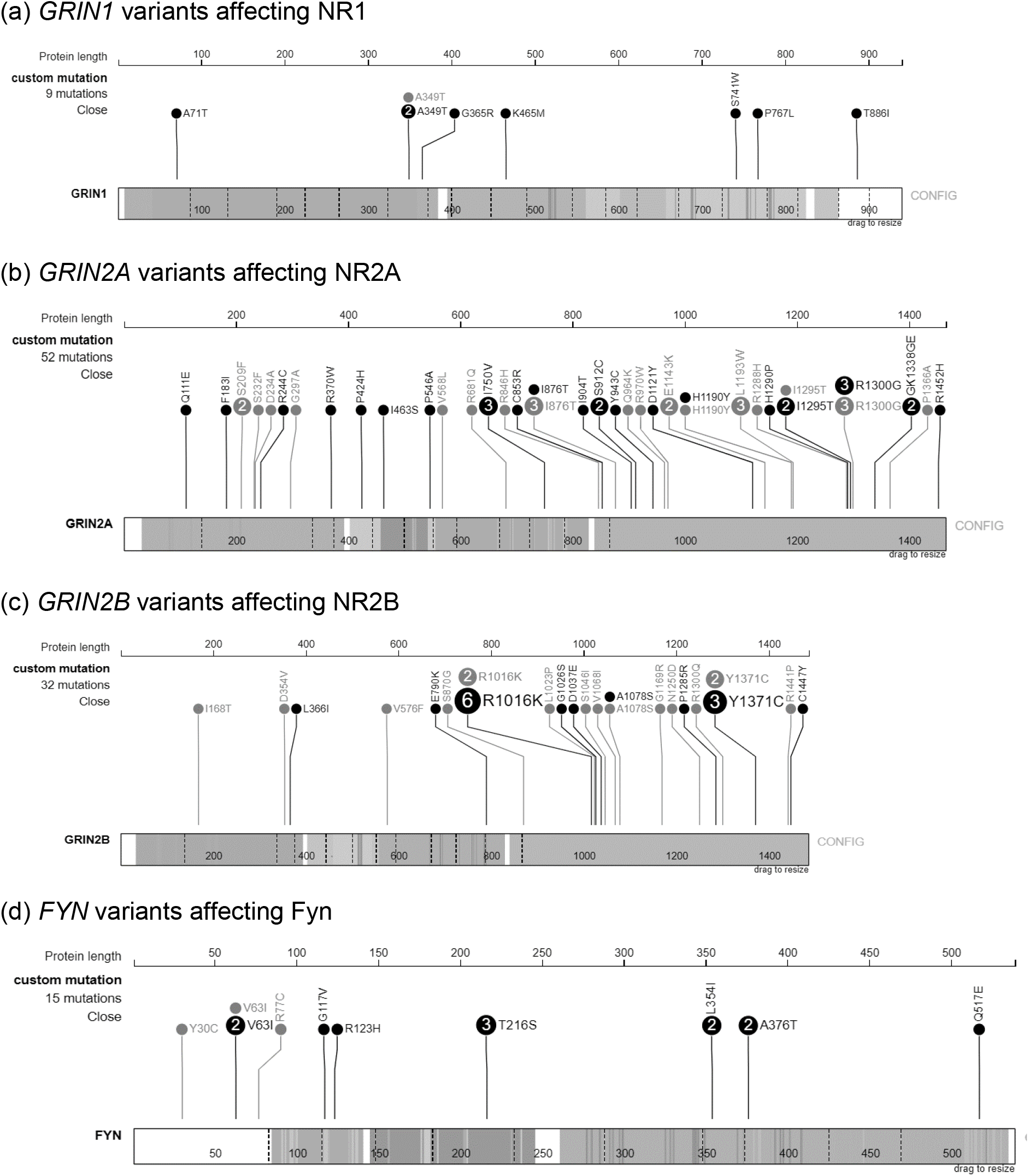
Variants from each gene in the relevant transcript as displayed by *ProteinPaint*. Variants seen in cases are coloured black while variants seen in controls are coloured grey. Where more than one case or one control has the same variant the number possessing it is shown within the circle and the size of the circle is increased proportionately.

With regard to phosphorylation sites, experimental support for each site can be quantified by the number of supporting references (high throughput / low throughput). The best supported site in NR2A is Y943 (134/2) and one case has a Y943C variant. In NR2B three cases and two controls have Y1371C but this site has only modest support (11/1). One control has a Y30C substitution in Fyn but this site has only low support (0/1). No other well supported sites were directly affected. However it may be worth noting that there two well supported sites adjacent to each other in Fyn, Y213 (1800/1) and Y214 (671/1), and that three cases have a T216S substitution. It is not inconceivable that this change two residues away might be sufficient to disrupt recognition of the site and hence impair phosphorylation.

In general, examination of the 3D localisation of the substitutions did not yield any new insights. However using model 2DQ7 (Kinoshita *et al*., 2006) for Fyn did reveal that A376T (observed in two cases) and Q517E (observed in one case) end up adjacent to each other in the folded protein. This is illustrated in Supplementary Figure 2.

On examining the distribution of variants across subjects in *GRIN1, GRIN2A, GRIN2B* and *FYN*, it emerged that one case had a damaging variant in *GRIN1* and a frameshift variant in *GRIN2A* while another subject had a damaging variant in *GRIN1* and in *GRIN2B*. No other subject had more than one variant in these genes.

## Discussion

The number of variants involved is too small for any definite conclusions to be reached. However there are a few points of interest. Firstly, there is no evidence to implicate disruption of serine-threonine phosphorylation or disruption of tyrosine phosphorylation by Src. Thus, if impaired tyrosine phosphorylation of NMDAR has any relevance to schizophrenia aetiology this seems likely to be specific to Fyn rather than Src. It is also not the case that amino acid substitutions in NMDAR preventing phosphorylation occur commonly in schizophrenia - we observe this in only a single case out of 4225. Nor is there strong evidence that variants interfering with Fyn autophosphorylation occur commonly. We see no definite examples of this and the closest we come to any support for this is the observation that three cases have an amino acid substitution two residues away from a tyrosine residue which is well supported as a phosphorylation site. We do observe that 3 cases and no controls have disrupting variants in GRIN2A and that 8 cases and 1 control have damaging variants in GRIN1. 14 cases and 3 controls have damaging variants in FYN but if we restrict attention to functional domains of the protein this applies to 10 cases and no control. Aside from that, we cannot discern any obvious pattern to the distribution of damaging variants. Overall, these possibly relevant variants were seen in about 0.5% of the cases.

These results are perfectly consistent with the hypothesis that the risk of schizophrenia may be increased by genetic variants impairing NMDAR functioning either directly or by reducing phosphorylation by Fyn. However this study provides at best only modest further support. It is hoped that the situation may become clearer as larger numbers of cases are sequenced. It might also be helpful to carry out functional studies to determine the effect of these variants *in vitro* and *in vivo*. This approach demonstrates ways in which additional information of possible biological relevance can be mined from sequencing studies.

## *In silico* investigation of coding variants potentially affecting the functioning of the glutamatergic N-methyl-D-aspartate receptor in schizophrenia

### Supplementary material

**Supplementary Figure 1.**
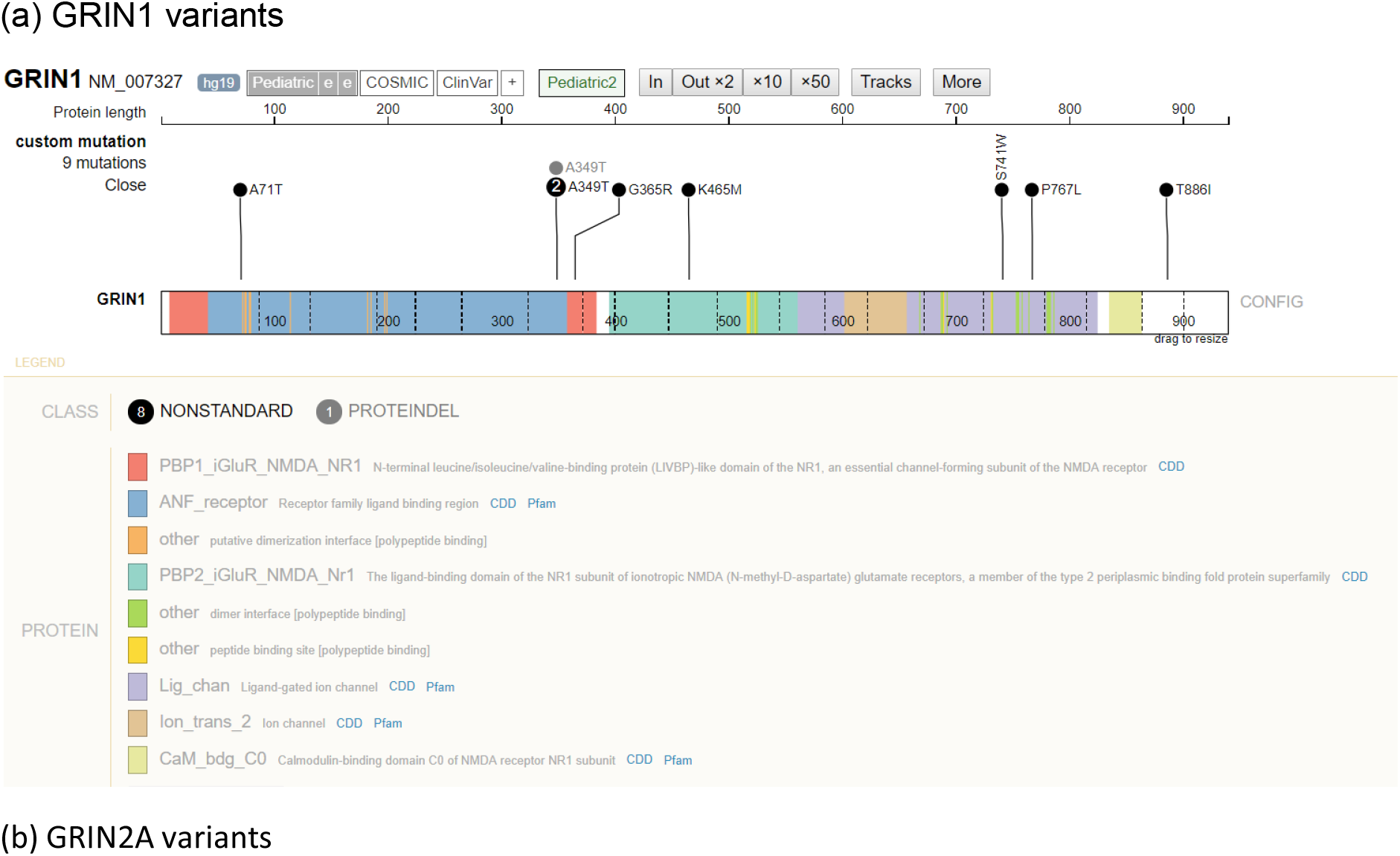
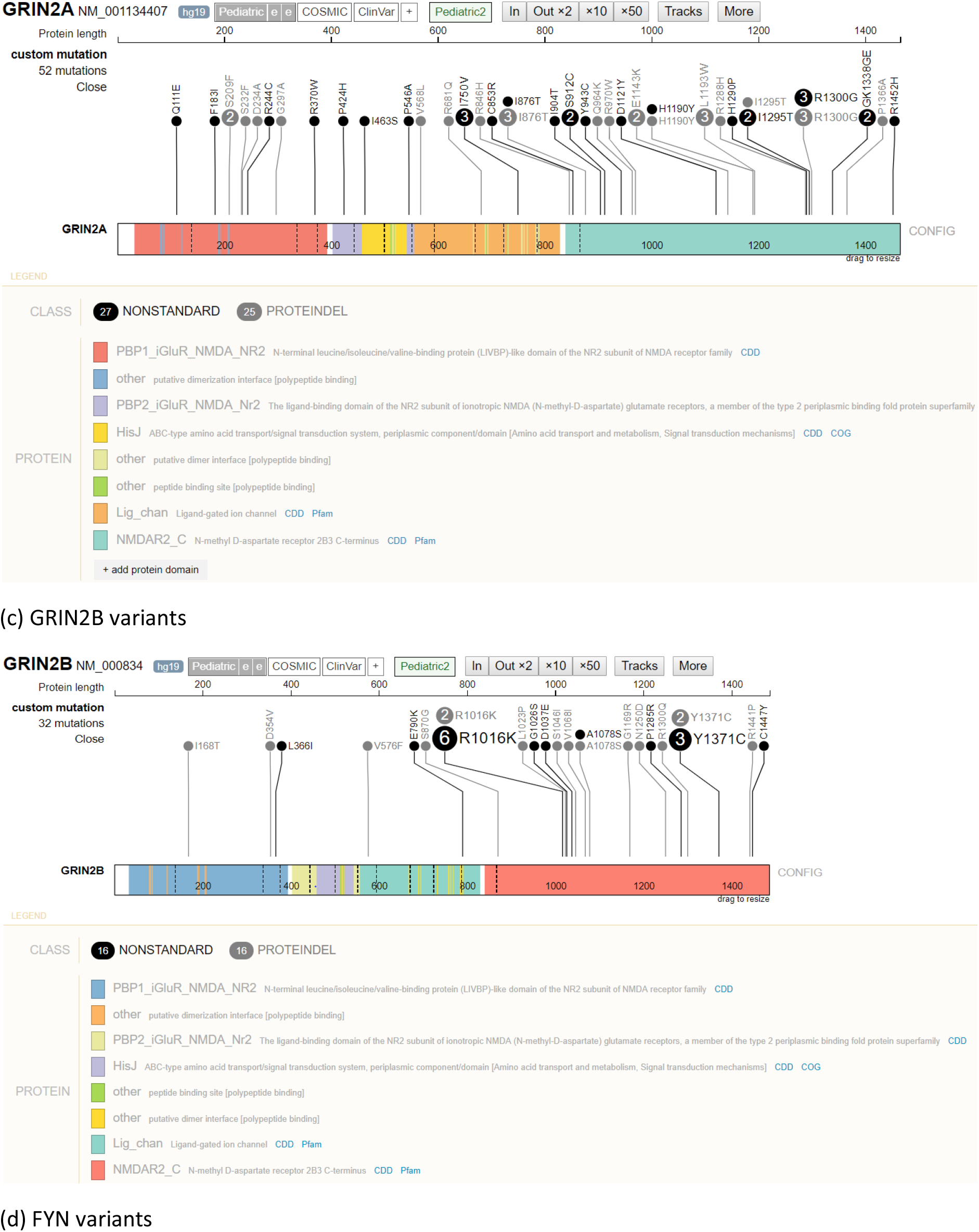
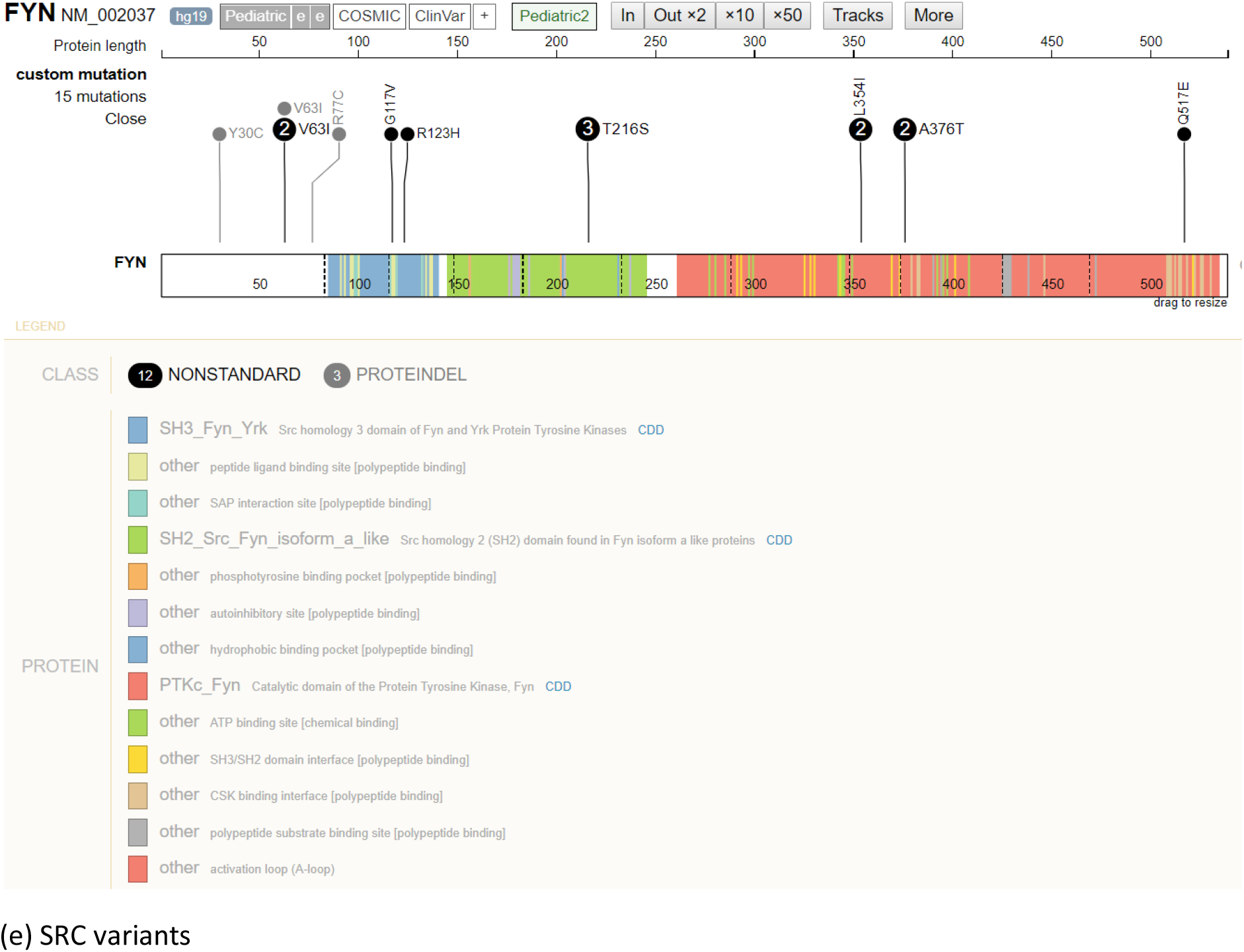
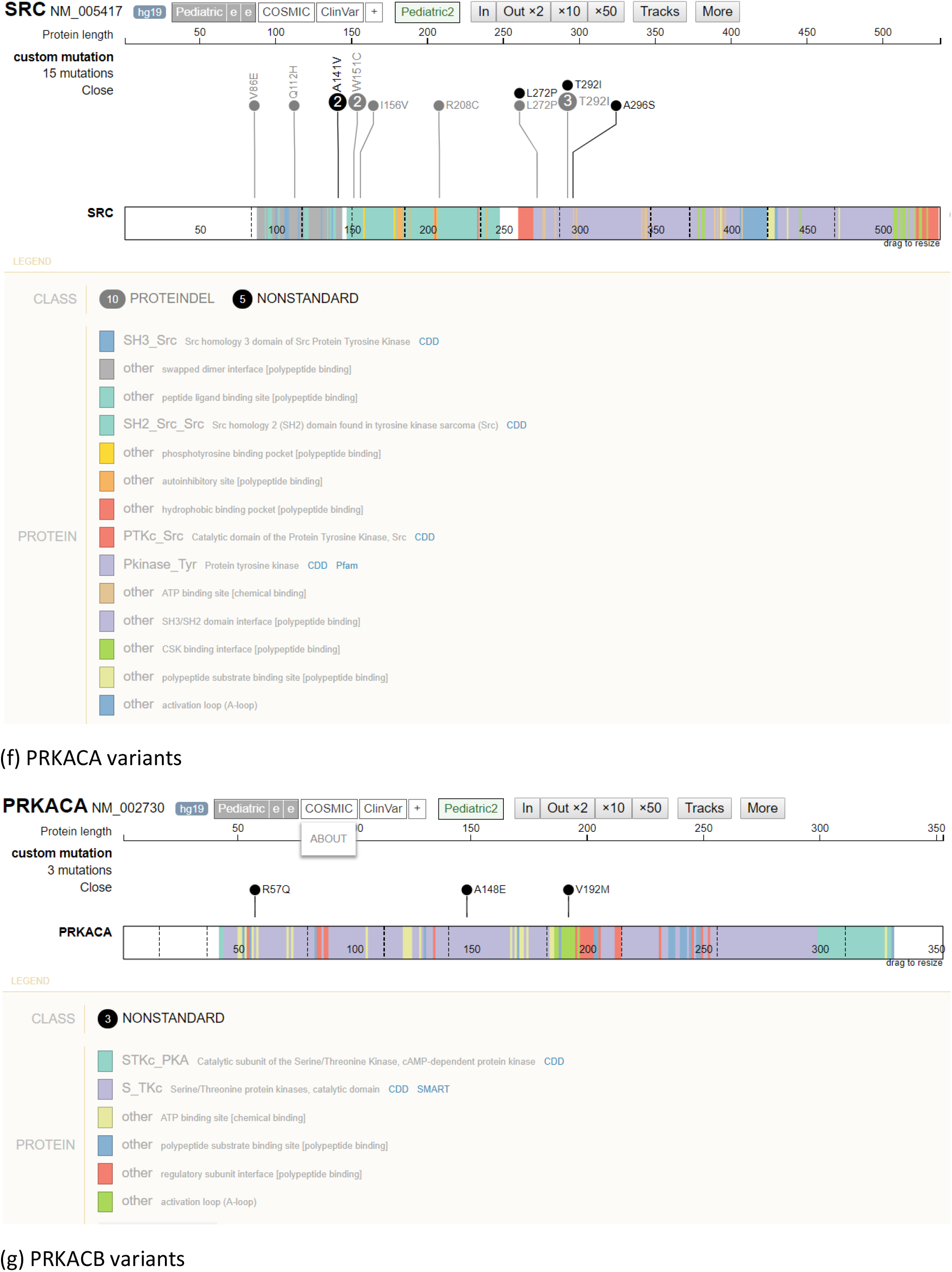
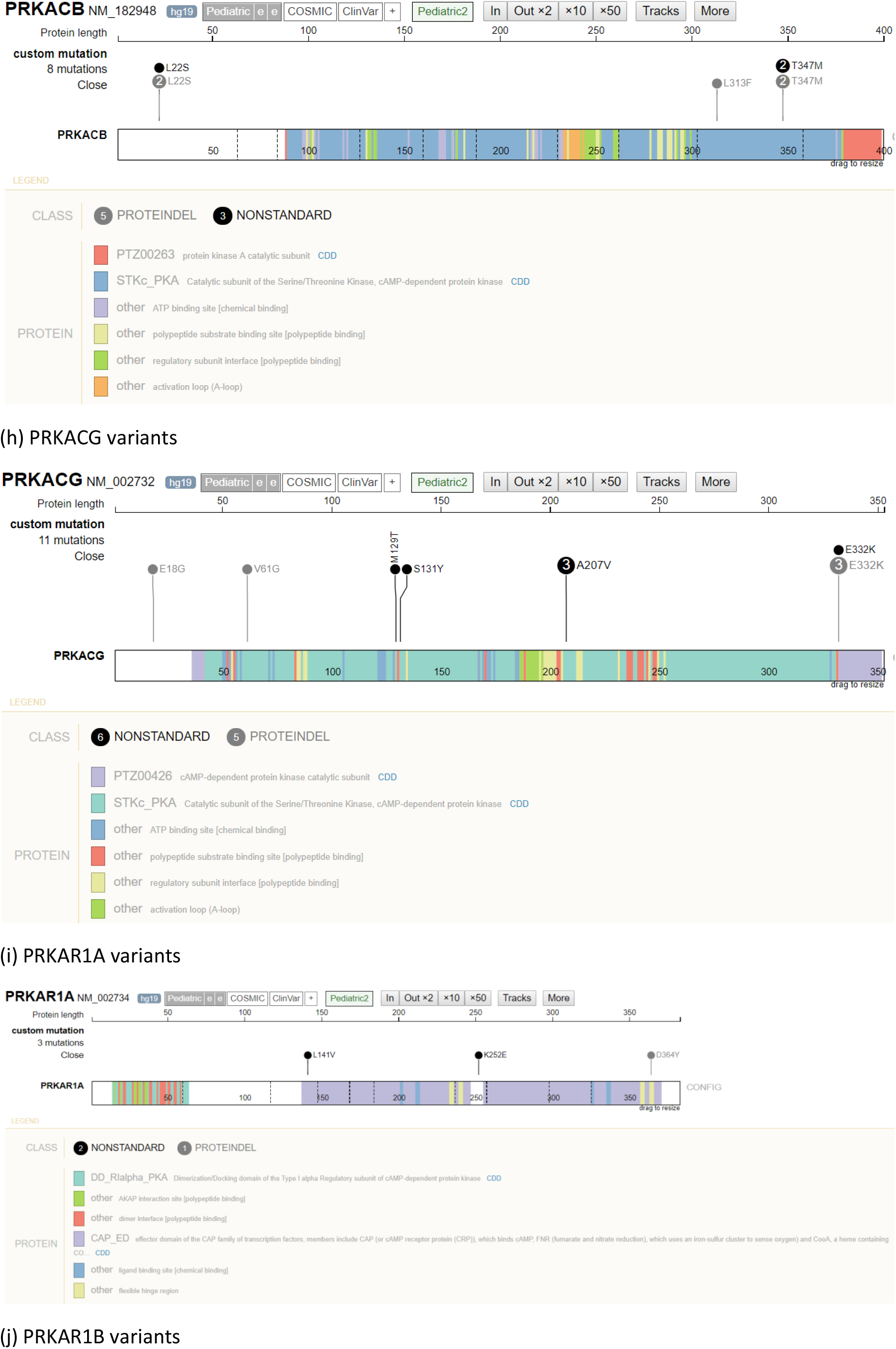
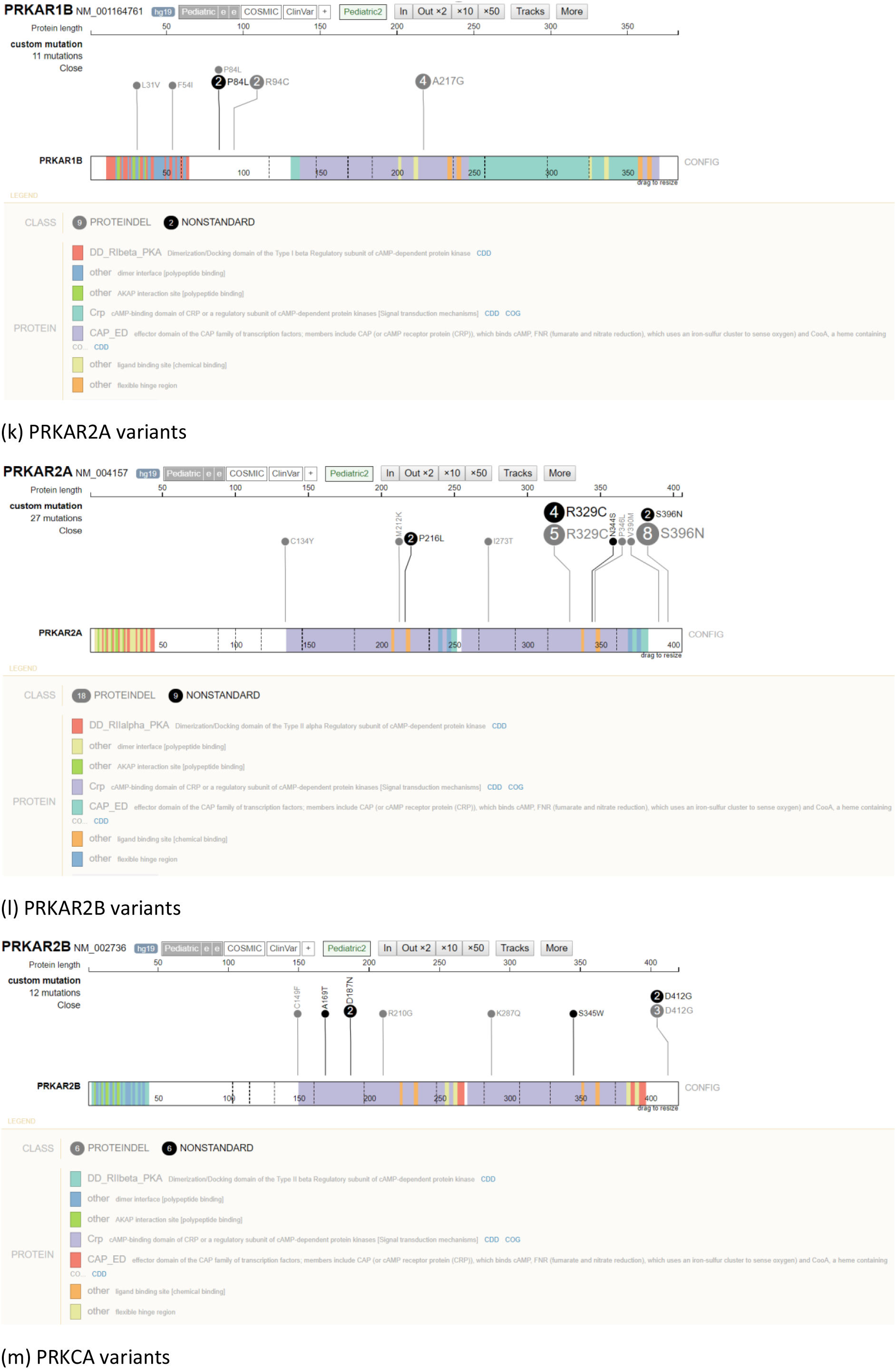
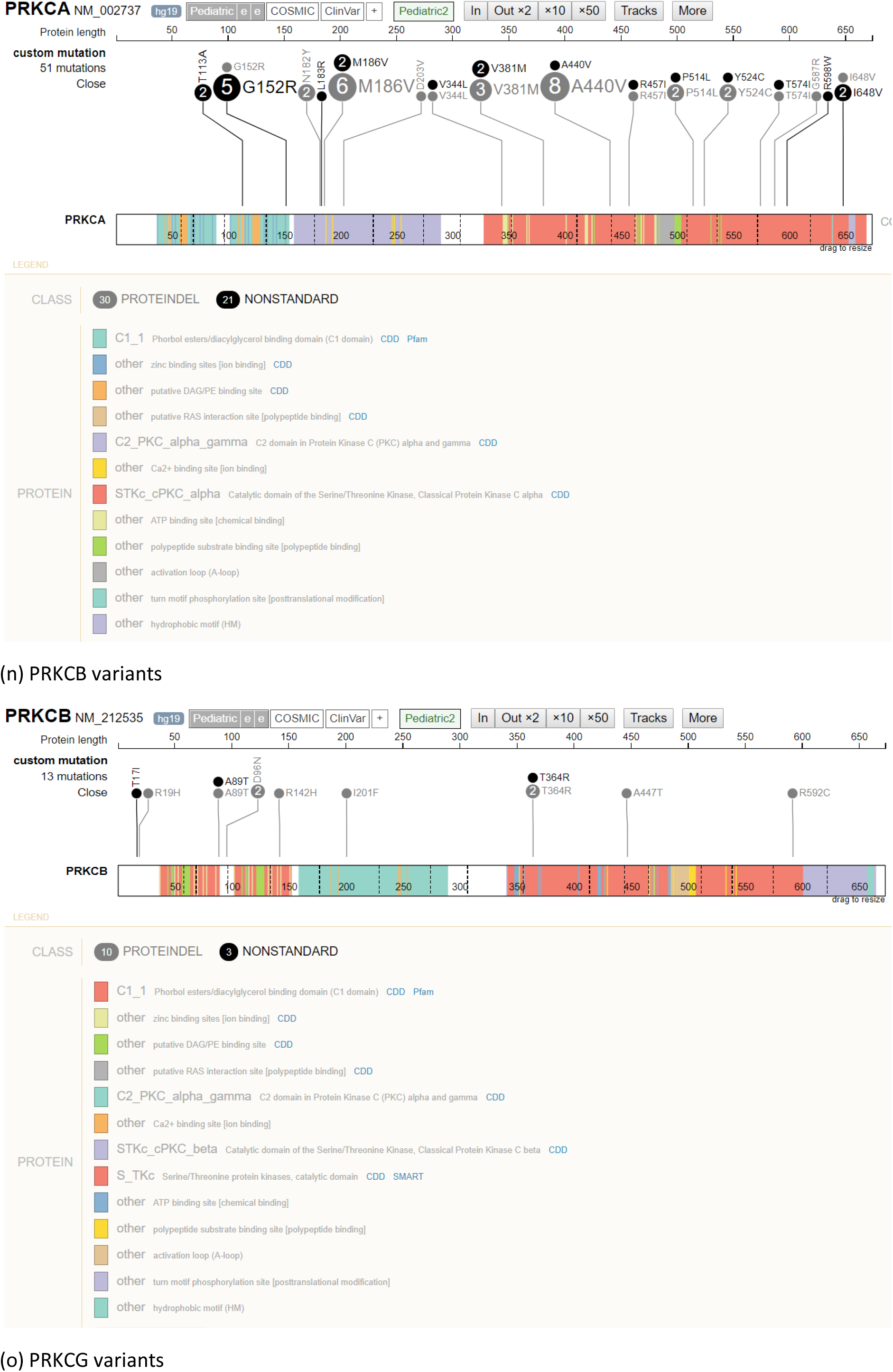
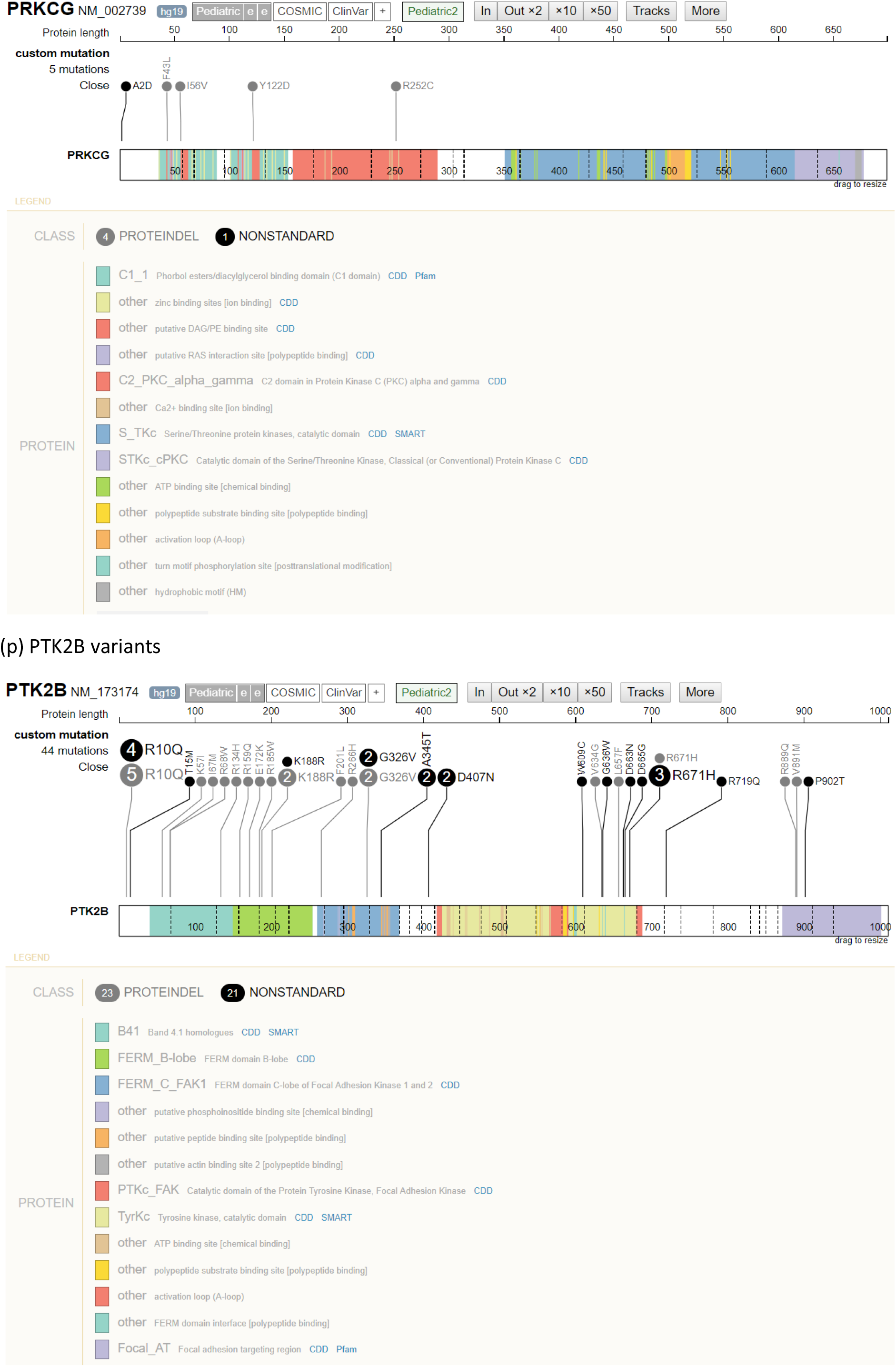
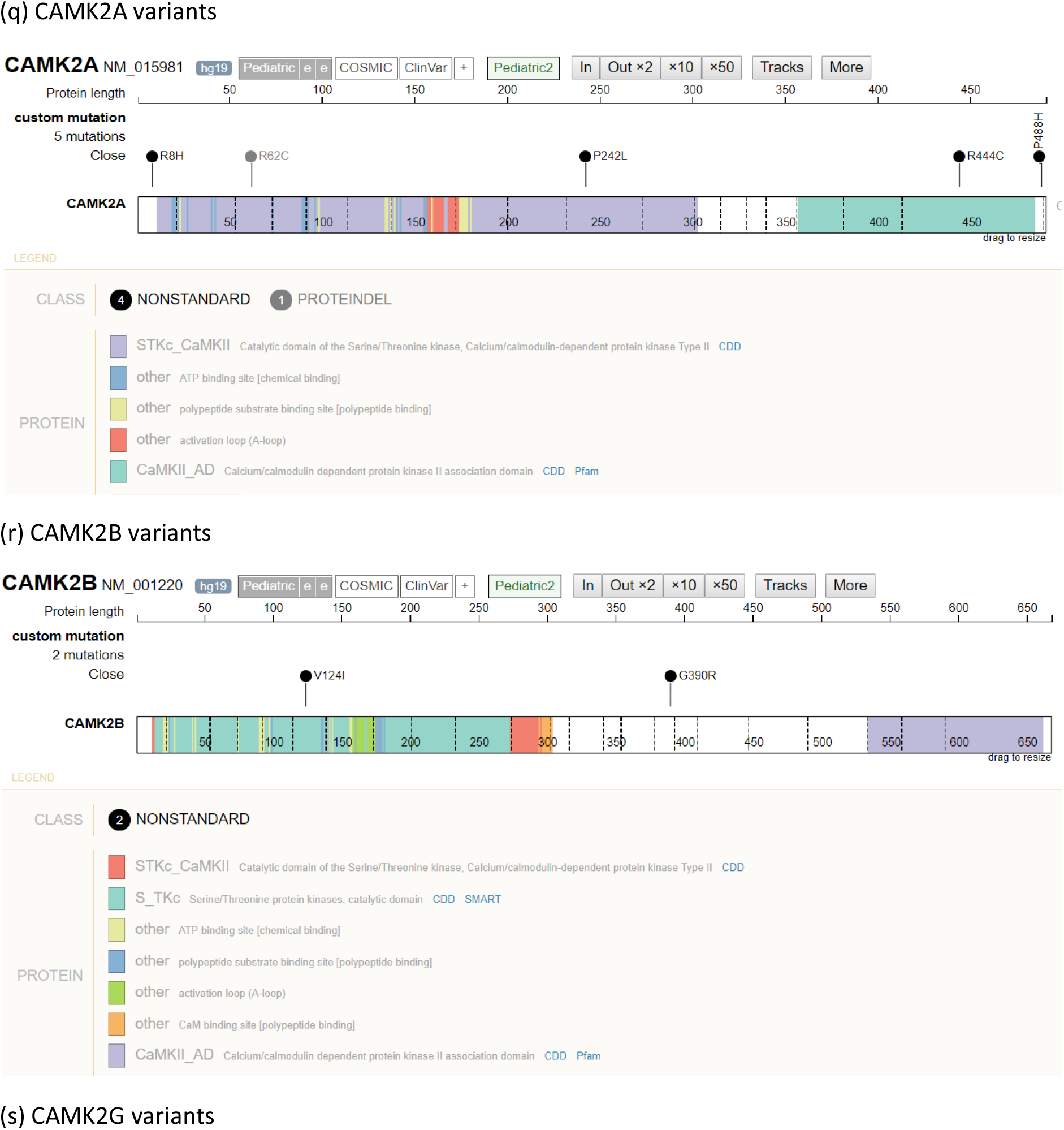
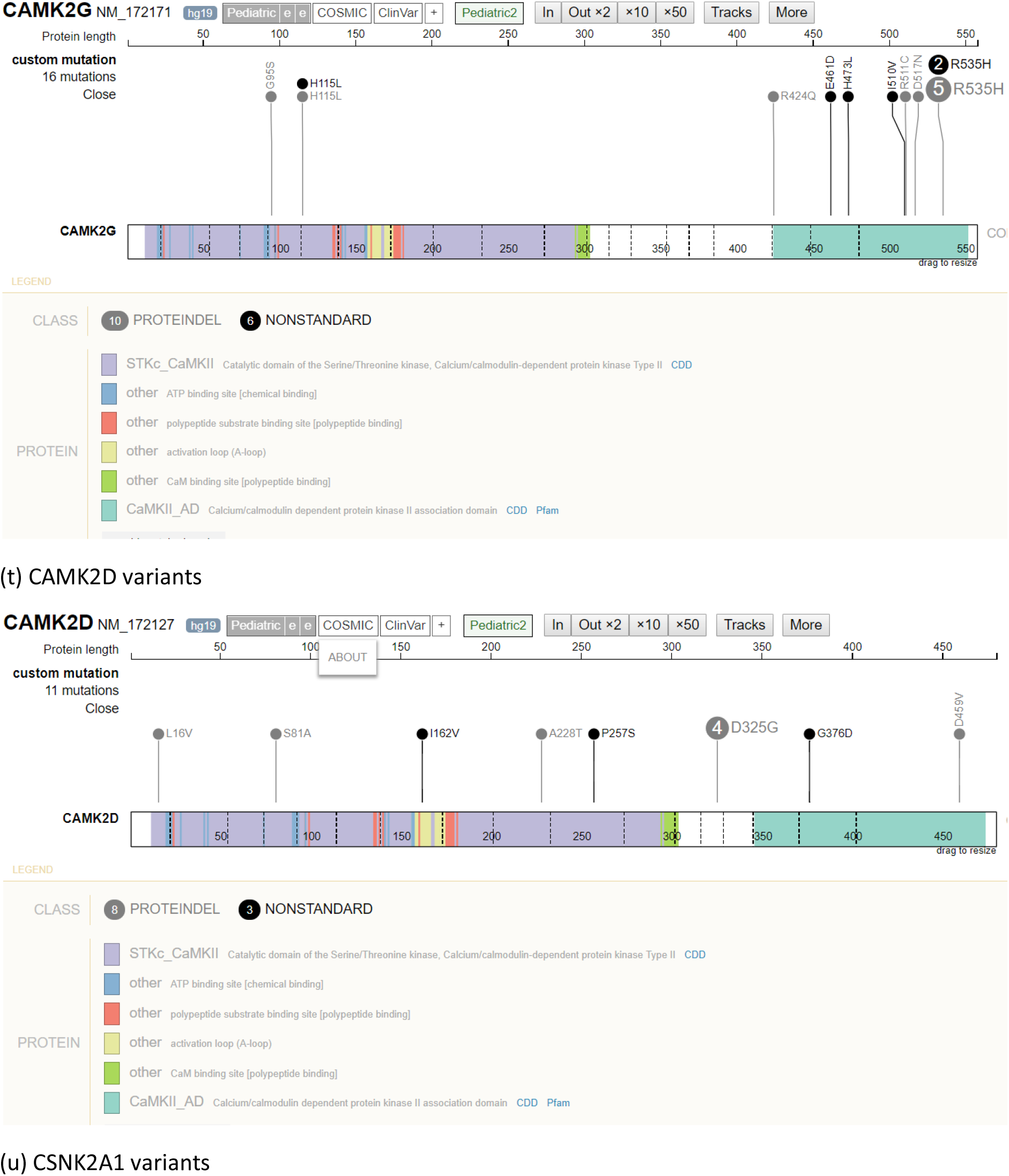
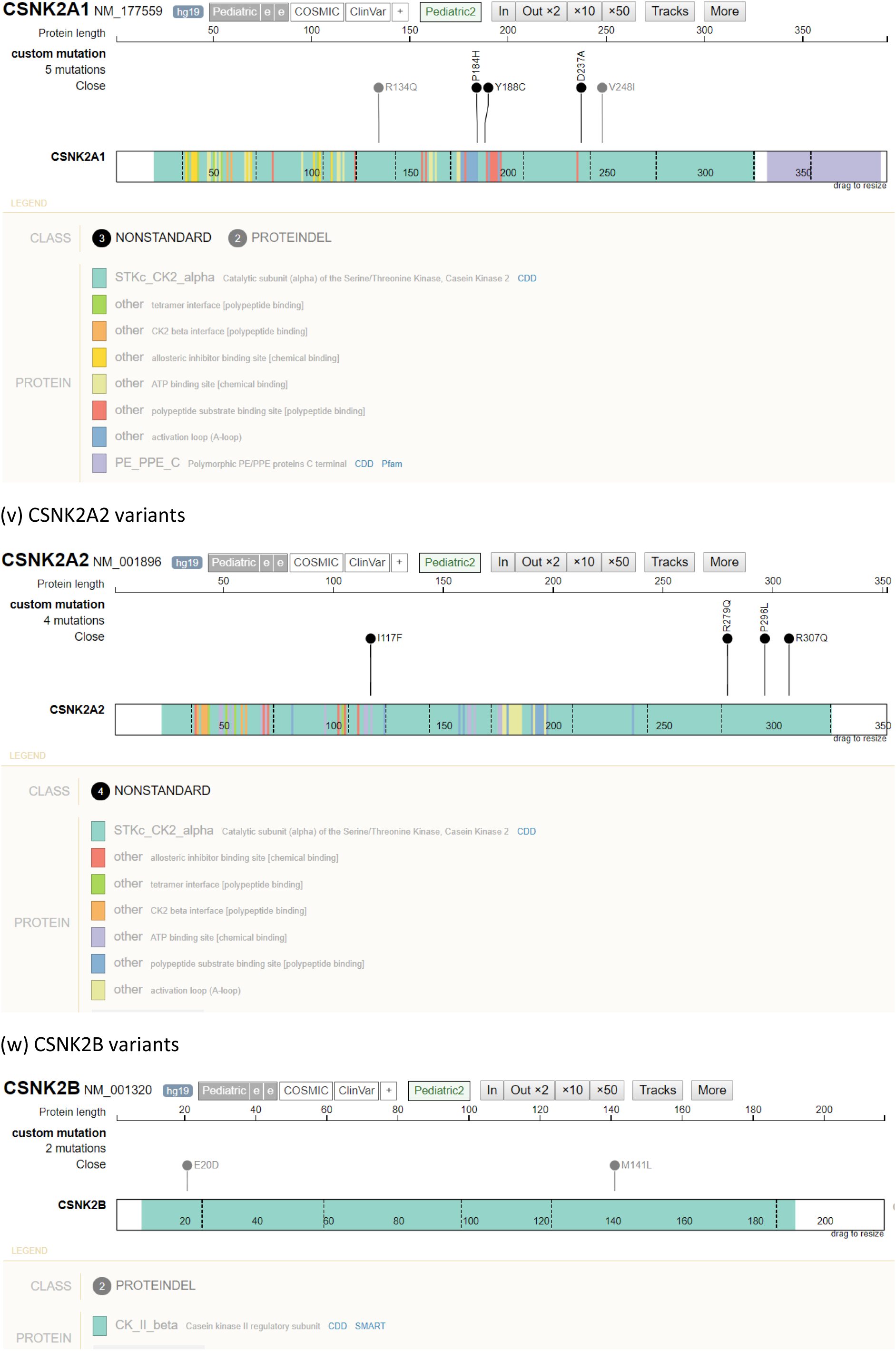
Variants from each gene in the relevant transcript as displayed by *ProteinPaint*. Variants seen in cases are coloured black and in the key labelled “NONSTANDARD” while variants seen in controls are coloured grey and labelled “PROTEINDEL”. Where more than one case or one control has the same variant the number possessing it is shown within the circle and the size of the circle is increased proportionately.

**Supplementary Figure 2.**
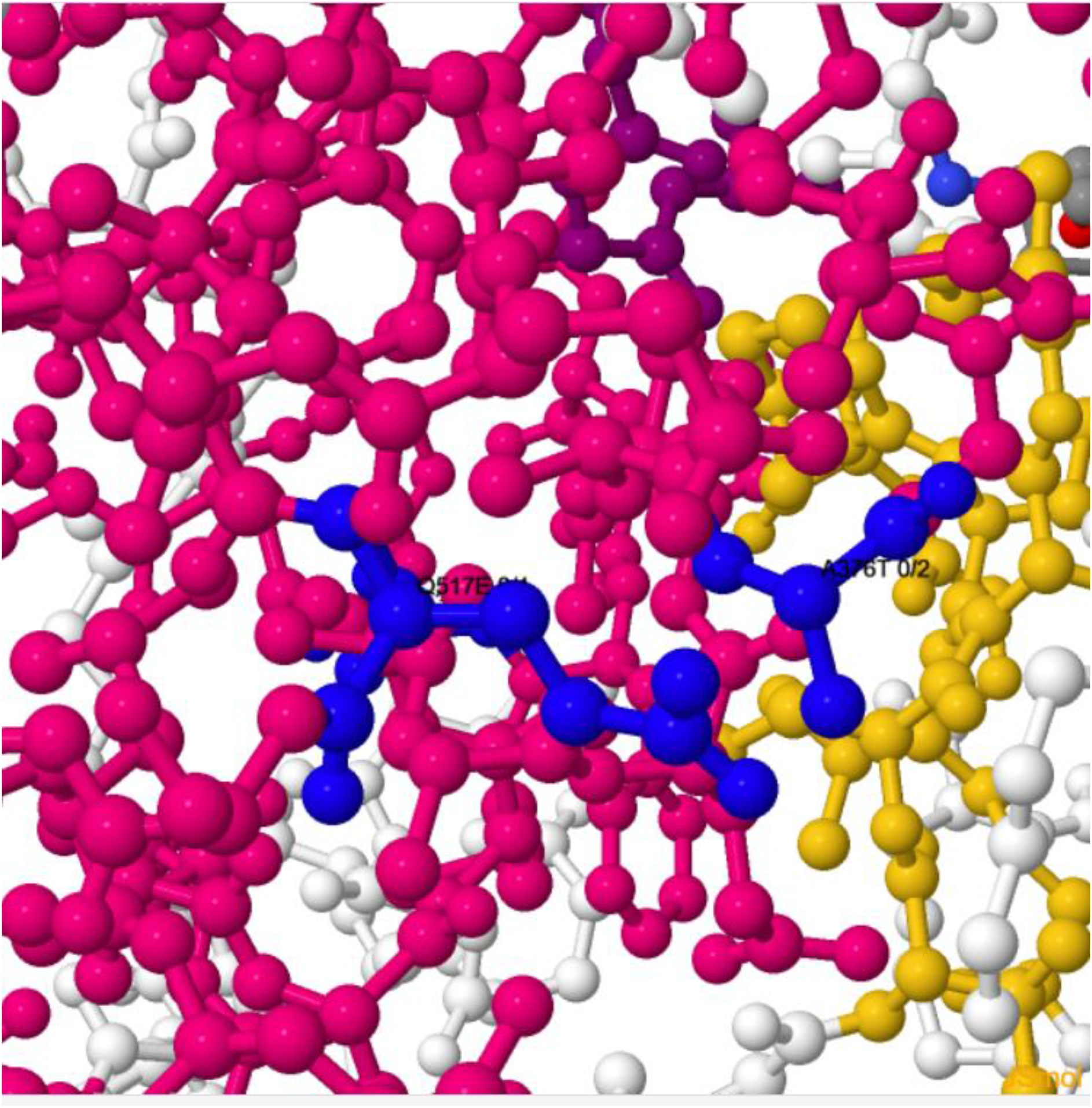
Screenshot from *JSmol* of structure 2DQ7 of Fyn showing that variants A376T and Q517E lie close together. Both are coloured blue with A376T (seen in two cases) on the right and Q517E (seen in 1 case) on the left.

